# Describing Chemical Kinetics in the Time Dimension with Mean Reaction Time

**DOI:** 10.1101/2022.02.01.478700

**Authors:** Chiwook Park

## Abstract

The chemical kinetics is such a fundamental topic in chemistry. By analyzing how the experimental conditions and parameters influence the reaction rate, chemists have deciphered the molecular details on how the chemical reactions occur. The quantitative analysis of the reaction rate requires the formulation of the rate equation, which describes the dependence of the reaction rate on the concentrations of the reactants and also the rate constants of the elementary steps. Though the methods to derive the rate equation from the kinetic model have been known for a century, it is still mathematically challenging to derive the rate equation for complex reactions with multiple steps, which requires a solution for simultaneous differential equations. Therefore, chemists frequently resort to the steady-state approximation or the numerical simulation. One way to avoid the mathematical difficulty is to describe the chemical kinetics in the time dimension. Describing the mean reaction time, the average of the time required for the completion of the chemical reaction, using the elementary rate constants of the kinetic model is much simpler mathematically than deriving the rate equation for the kinetic model. Here, we describe the basic rules for the derivation of the formula of the mean reaction time, derive the generalized equation for the mean reaction time, and analyze the components of the formula to determine how the individual steps in the complex reaction contribute to the mean reaction time. Being the ensemble-averaged value, the mean reaction time does not provide the information on the actual distribution of the reaction time of individual chemical entity. However, the formula of the mean reaction time reveals invaluable insights on how the energy levels of the ground state and the transition states affect the kinetics of the complex reaction and offers a way to identify the most time-consuming process of the complex reaction in a straightforward manner. We also apply the mean reaction time to enzyme kinetics and demonstrate that one can derive the expressions of the kinetic parameters (*k*_cat_/*K*_M_ and *k*_cat_) in a surprisingly simple way even without resorting to the steady-state application.

---

When a chemical reaction occurs in multiple steps, the rate equation is expressed as a function of elementary rate constants of the steps. Understanding how the elementary rate constants affect the overall reaction rate is critical in analyzing the kinetic data and deciphering the kinetic mechanism of the reaction. However, the derivation of the rate equation for the complex reaction is mathematically challenging, even for a chemical reaction with only a few steps, because the rate equation is in principle a solution of simultaneous differential equations. Scheme 1 shows a kinetic model for a simple enzyme-catalyzed reaction that occurs in three steps between one substrate (S) and one product (P):

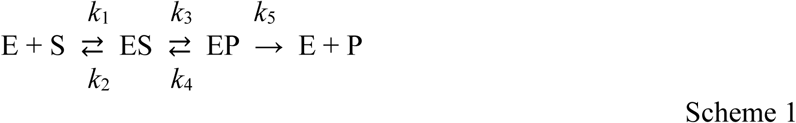

Derivation of the rate equation requires solving the following four simultaneous differential equations:

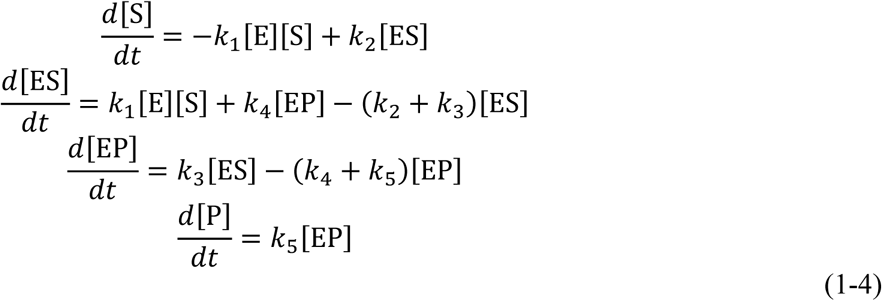

In some cases, one can obtain the analytical solution using the Laplace transform or the determinant methods.^1^ But, the resulting equation is usually extremely complex and offers little insight on the kinetic mechanism. Due to this difficulty, chemists typically employ various approximations, such as steady-state approximation or rapid equilibria approximation, and convert the simultaneous differential equations to the simultaneous linear equations.^1, 2^ When the steady-state approximation is employed for the above case,^3, 4^ the rate equation is derived to be:

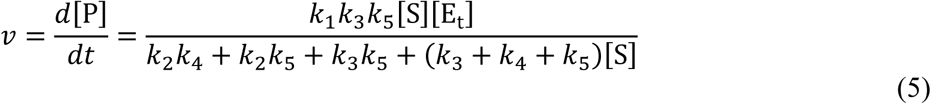

which is in the form of the Michaelis–Menten equation. The two fundamental kinetic parameters of the enzyme kinetics, *k*_cat_ and *k*_cat_/*K*_M_ are then expressed as functions of the elementary rate constants:

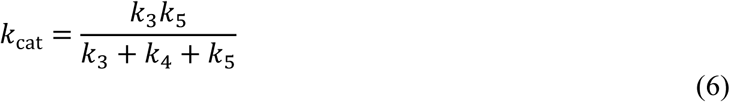

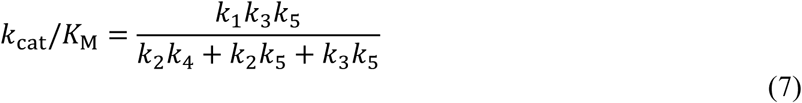

Deciphering from the above equations how individual steps in the kinetic model contributes to the two kinetic parameters is still not so intuitive.

Here we report a radically different approach to describe chemical kinetics using the mean reaction time (τ_rxn_), which is the average time required for the completion of the reaction, for complex reactions comprised of first-order elementary reactions. While the first-order rate constant has the unit of the inverse of time, the mean reaction time has the unit of time, which gives more intuitive appreciation of the speed of the reaction in the real physical dimension. Moreover, this approach does not require solving differential equations or any mathematical approximations. Based on a few simple rules, one can easily derive the expression for the mean reaction time with the elementary rate constants of the kinetic model. In case of the reaction only with irreversible steps, the mean reaction time is simply the sum of the mean reaction times of the individual steps. For the reaction with reversible steps, the extra reaction time needs to be added due to the repeated forward and reverse reactions at the reversible steps. We report here a step-by-step approach to calculate the extra reaction time required for the reversibility. The expression of the mean reaction time as the sum of several terms, which also have the unit of time, allows us to examine in an intuitive way how individual steps contribute to the mean reaction time. Finally, we demonstrate the feasibility of the use of the mean reaction time by deriving the expression of the mean reaction time for the enzyme-catalyzed reaction in Scheme 1 (and the expressions for *k*_cat_ and *k*_cat_/*K*_M_) even without using the steady-state approximation.

## Mean reaction time

The mean reaction time (τ_rxn_) of a reaction is defined as the average of the time required for the reactant to be converted to the product. When the reaction starts with only with the reactant and with none of the product, the mean reaction time is the statistical mean of the reaction time weighted by the population of the product (P) formed at each time point.

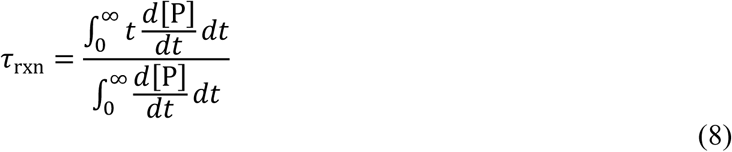

For the first-order elementary reaction shown in Scheme 2,

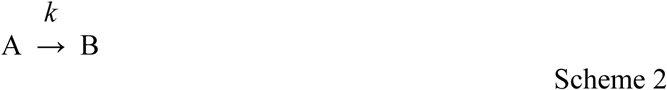

the kinetic equation for the production of B is a single-exponential function as:

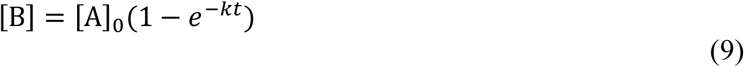

where [A]_0_ is the concentration of A at *t* = 0. When calculated with Eq. 8, the mean reaction time of this reaction is 1/*k*.

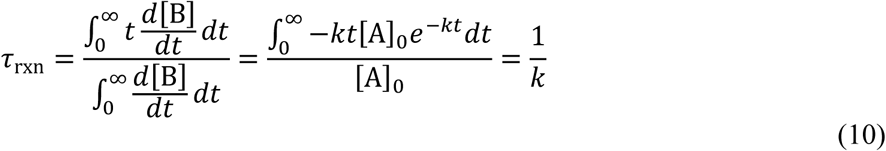

For the first-order elementary reaction, the mean reaction time (1/*k*) is also called the time constant (τ), the residence time, or the mean lifetime of A. The mean reaction time of the first-order reaction is greater than the half-life of the reaction, which is the median of the reaction time (ln2/*k* ∼ 0.693τ) (Figure 1).

**Figure 1.**
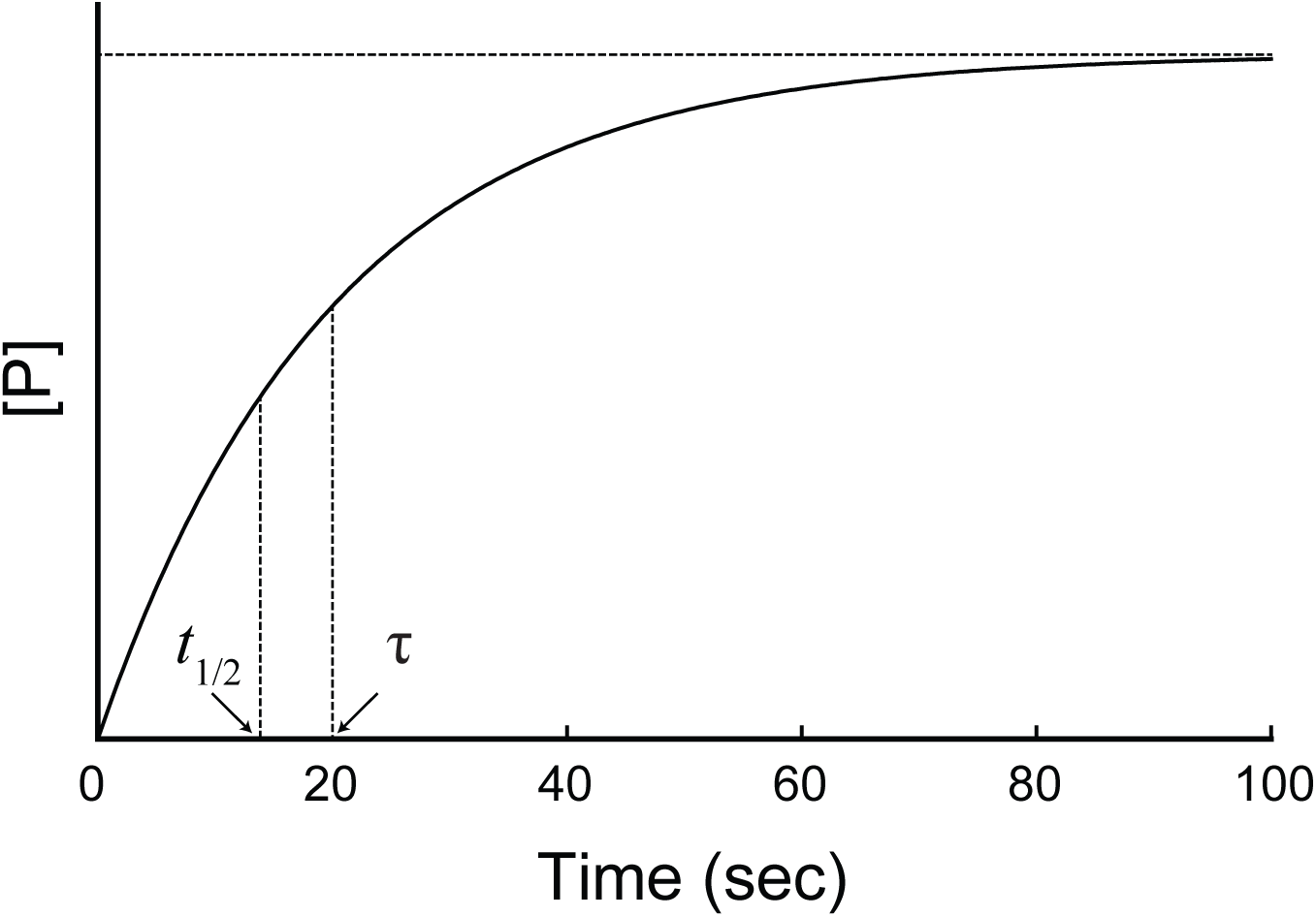
Comparison of the time constant (τ) and the half-life (*t*_/2_) of a first-order reaction. The production of P is simulated using the first-order rate constant of 0.05 s^-1^. The time constant of the reaction (20 sec) is the mean of the reaction time, and the half-life of the reaction (13.9 sec) is the median of the reaction time.

When the reaction is a complex reaction with multiple elementary steps, the kinetic equation for the production of the product may or may not be a single-exponential function. If the kinetic equation is a single-exponential function, the rate constant for the overall reaction would be equal to 1/τ_rxn_ as shown in Eq. 10. We define τ_rxn_ in this case as being “explicit”; we can reconstruct the kinetic equation like Eq. 9 from τ_rxn_. If the kinetic equation is not a single-exponential function, the mean reaction time is still defined as in Eq. 8. However, a rate constant corresponding to 1/τ_rxn_ would not exist, and we cannot reconstruct the kinetic equation. We define τ_rxn_ in this case as being “implicit”. Like rate constants, the mean reaction time is independent of the concentration of the reactant. The conversion of each reactant to the product is an independent event. Each reactant has the same probability of the conversion at each step, regardless of the quantity of the reactant. Therefore, the mean reaction time is a characteristic constant of the reaction under the given experimental condition.

## Mean reaction time of two-step reactions

To calculate the mean reaction time for a complex reaction, we sum up the mean lifetime of the reactant or the intermediate in the reaction trajectory until the reaction is complete. We perform the calculation based on the following two simple rules on the mean reaction time of the reactions with two steps in sequential and parallel.

### Rule 1

When a reaction occurs in two sequential irreversible steps as shown in Scheme 3, the mean reaction time (τ_rxn_) is the sum of the reciprocals of the two kinetic constants.

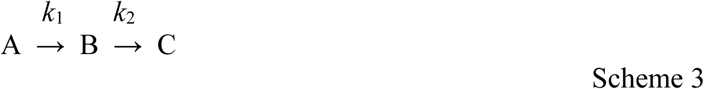

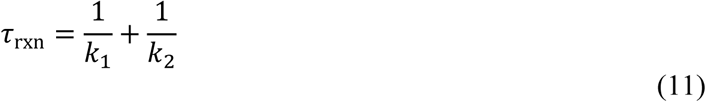

Obviously, 1/*k*_1_ and 1/*k*_2_ are the mean lifetimes of A and B, respectively. As long as the two steps are independent to each other, which means that B has no memory of when it has formed from A, the mean reaction time is the sum of the mean lifetime of A and B. The half-lives of A and B do not have this additive property because the half-life is the median, not the mean of the population. We can prove Eq. 11 by deriving the kinetic equation for the production of C according to the mass action law.

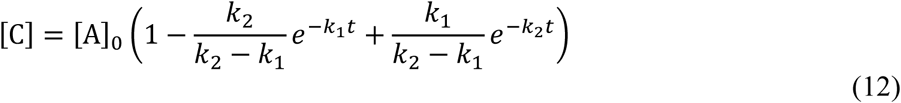

Here we assume that *k*_1_ is not equal to *k*_2_. When we calculate the mean reaction time using Eqs. 8 and 12, we obtain the same expression for the mean reaction time as Eq.11.

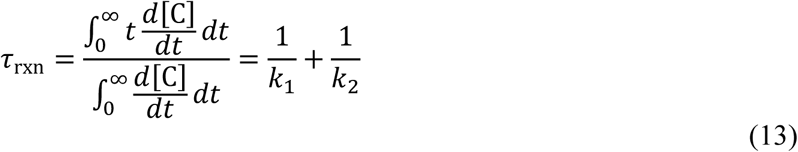

This rule is valid with reactions with more than two sequential irreversible steps. The mean reaction time is sum of the reciprocals of all the first-order rate constants of the steps in the reaction.

In general, τ_rxn_ for the reaction with two sequential irreversible steps is implicit; no single rate constant corresponding to 1/τ_rxn_ exists. When one step is significantly slower than the other, however, the slower step becomes the rate-limiting step, and τ_rxn_ becomes explicit; a single rate constant corresponding to 1/τ_rxn_ exists. When *k*_1_ << *k*_2_, the mean reaction time is determined dominantly by *k*_1_, which is 1/τ_rxn_. This is the case where the steady-state approximation is applicable, and the rates for both production of C and consumption of A are expressed with a single-exponential function with *k*_1_ as the rate constant. When *k*_1_ >> *k*_2_, the mean reaction time is determined dominantly by *k*_2_, which is 1/τ_rxn_. In this case, A converts to B completely first, and both consumption of B and production of C are expressed with a single-exponential function with *k*_2_ as the rate constant.

### Rule 2

When a reaction occurs in two parallel irreversible steps as shown in Scheme 4, the mean reaction time (τ_rxn_) is the reciprocal of the sum of the two kinetic constants.

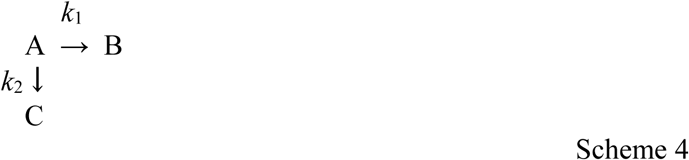

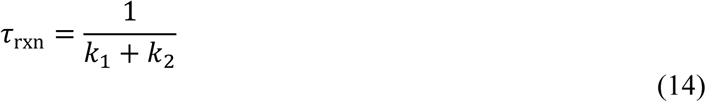

The reactant (A) disappears through the two parallel steps, and the overall reaction is still first-order with the rate constant of *k*_1_ + *k*_2_. Therefore, the mean lifetime of A is 1/(*k*_1_+*k*_2_). The final concentrations of B and C are determined by the ratio of the rate constant for their production to *k*_1_ + *k*_2_.

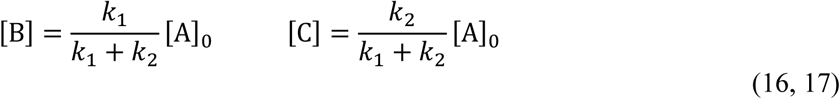

This rule is valid for the reaction with more than two parallel steps. The mean reaction time is the reciprocal of the sum of all the first-order rate constants, and the partition of A to each product is proportional to the rate constant for each step.

## Mean reaction time of the complex reaction

The calculation of the mean reaction time of the complex reaction with only irreversible steps is trivial as discussed above. However, when the reaction contains one or more reversible steps, the reaction trajectories become complex because some fraction of the reaction trajectories may cross the same step more than once. Scheme 5 shows a simple case in which a reversible step is followed by an irreversible step:

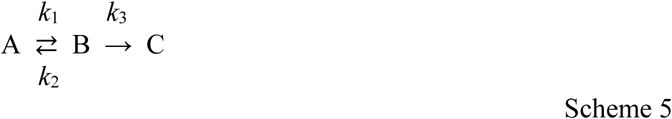

When B forms from A, B has two fates. Some of B proceeds to C and finishes the reaction, but the rest of B returns to A and start the whole process again. The two fates of B can be treated as two parallel steps discussed above in Rule 2.

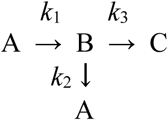

We can calculate the partitions of B into the two steps using Eqs. 16 and 17. Here, we define the partition of B that proceeds to C as *p*.

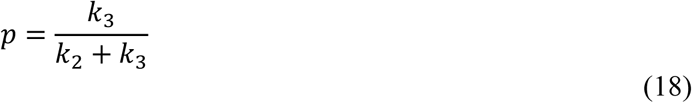

Then, the partition of B that returns to A is 1–*p*. According to Rule 2, the mean lifetime of B is 1/(*k*_1_+*k*_2_). The mean reaction time is calculated by summing the reaction time for each fraction that finishes the reaction on a probability tree diagram (Figure 2A). This problem is similar to the well-known drunkard’s walk problem^5^ with home (or a cliff) at the end. Different from the standard drunkard’s walk problem, the lifetime and the partition at each location varies. The sum of the reaction time for each fraction that finishes the reaction (fractions underneath C in Figure 2A) weighted by the size of the fraction is expressed as a sum of an infinite series:

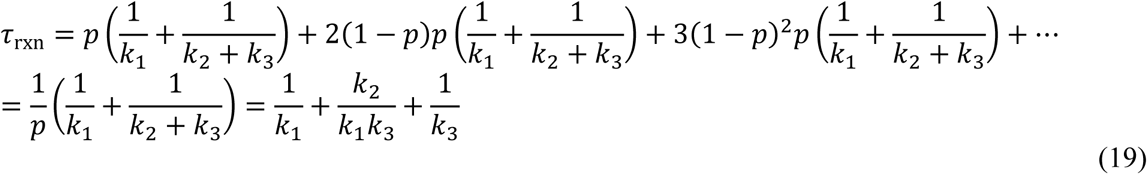

**Figure 2.**
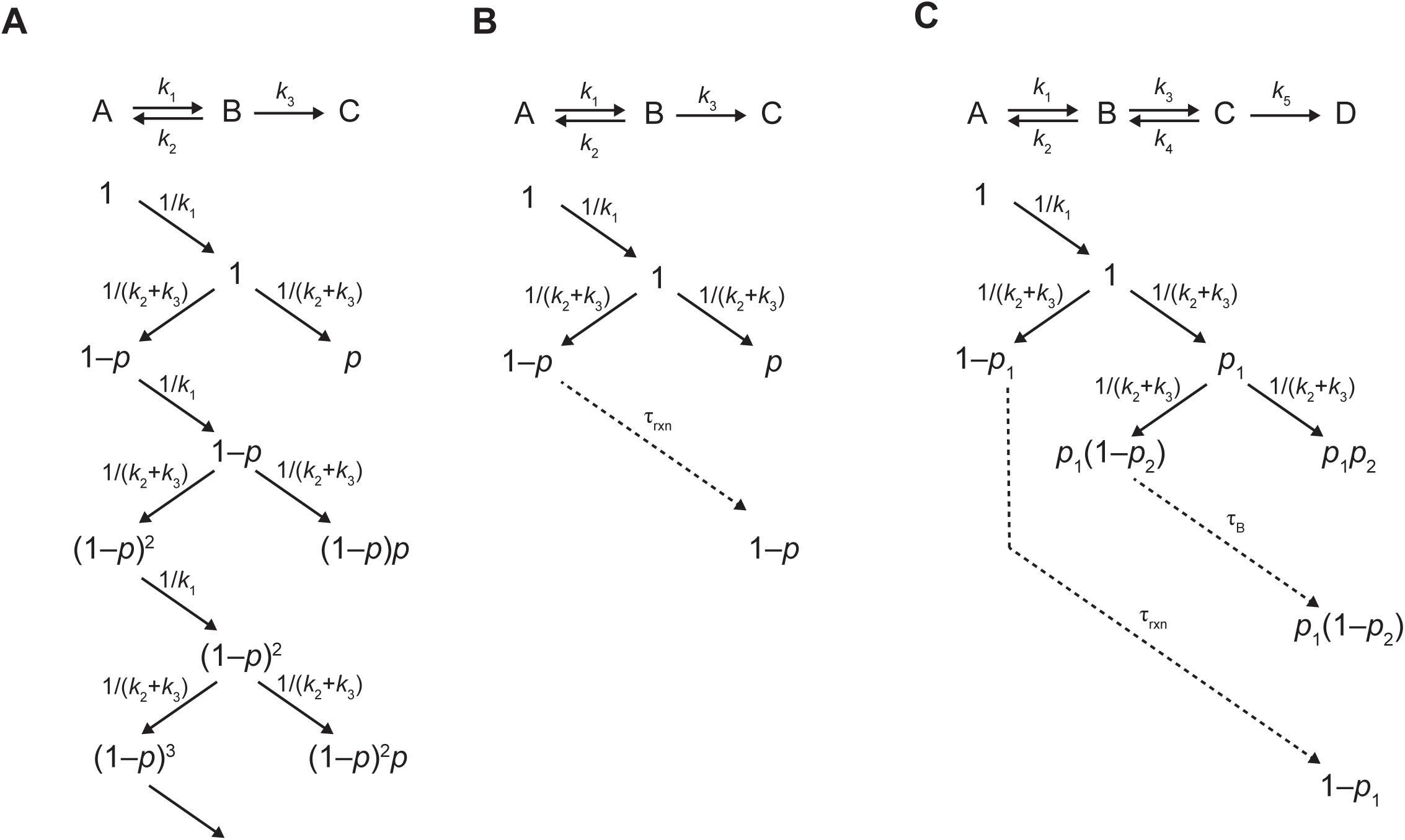
Probability tree diagrams for the mean reaction time. **(A)** The probability tree diagram for the reaction in Scheme 5. The partition of B that proceeds to C is defined as *p* (Eq. 18). The residence time added at each step is indicated above the arrows. The mean reaction time is the sum of the reaction time for each fraction that finishes the reaction weighted by the fraction (underneath of C, the final product) (Eq. 19) **(B)** The probability tree diagram for the reaction in Scheme 5 simplified with the use of τ_rxn_. **(C)** The probability tree diagram for the reaction in Scheme 6 simplified with the use of τ_rxn_. The partitions of B that proceeds to C and C that proceeds to D are defined as *p*_1_ and *p*_2_, respectively (Eqs. 24 and 25). τ_B_ is the mean reaction time required for B to finish the reaction, and τ_rxn_ is same as 1/*k*_1_ + τ_B_ (Eq. 27).

Compared with the mean reaction time for the reaction with only irreversible steps (Scheme 3), the mean reaction time for this reaction has one extra term, *k*_2_/*k*_1_*k*_3_. When *k*_2_ is set to 0, which means the first step becomes irreversible, the extra term disappears. The mean reaction time is increased by *k*_2_/*k*_1_*k*_3_ because of the reversibility of the first step. Therefore, *k*_2_/*k*_1_*k*_3_ is the kinetic cost of the reversibility that allows B to return to A. The extra time imposed by the reversible step is indeed the increase in the total residence time of A, which is the sum of the mean lifetime of the initial population of A and that of the fractions that return to A. Using the probability tree diagram (Figure 2A), we determine the total residence time of A:

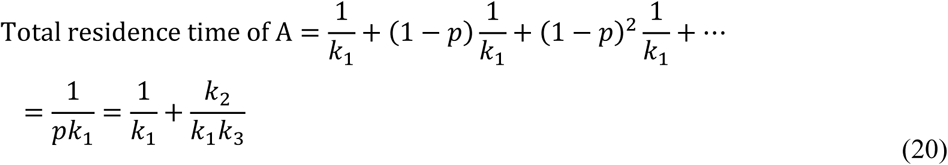

which shows that the total residence time of A is increased by *k*_2_/*k*_1_*k*_3_ due to the returning fractions. This result also suggests that the total residence time of B is not affected by the reversibility of the first step. Using the same approach, we determine the total residence time of B:

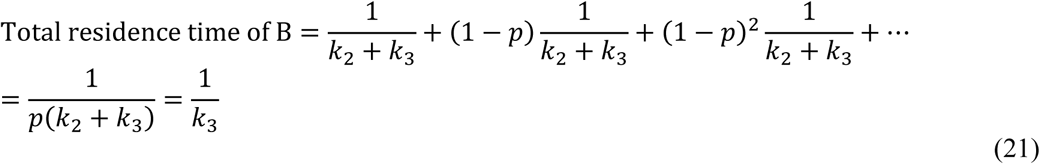

which is identical to the mean lifetime of B when the first step is irreversible. Obviously, the sum of the total residence times of A (Eq. 20) and B (Eq. 21) is the mean reaction time of the reaction (Eq. 19).

An even simpler way to calculate the mean reaction time is the recursive use of τ_rxn_ (Figure 2B). After the first trip from A to B, the fraction of *p* proceeds to C and finishes the reaction, and the mean reaction time for this fraction is 1/*k*_1_ + 1/(*k*_1_+*k*_2_). The fraction of 1–*p* returns to A and starts the reaction from the beginning and requires τ_rxn_ again to finish the reaction. The mean reaction time for this fraction is, therefore, 1/*k*_1_ + 1/(*k*_1_+*k*_2_) + τ_rxn_. The mean reaction time is then expressed as:

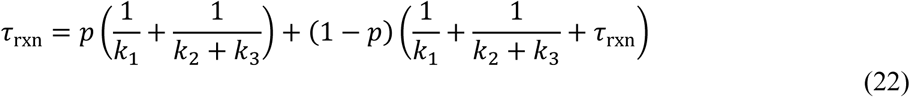

By solving Eq. 22 for τ_rxn_, we obtain the same expression for τ_rxn_ as in Eq. 19:

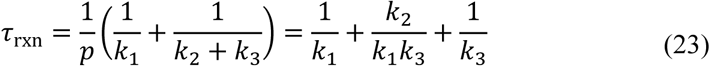

Now we determine the mean reaction time for the reactions with more than one reversible step. The reaction in Scheme 6 has two reversible steps followed by one irreversible step:

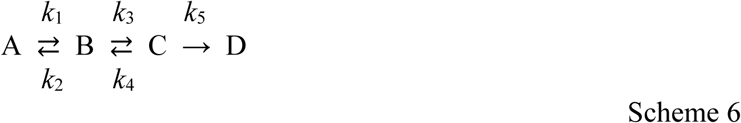

As B and C have two fates of proceeding and returning, the reaction in Scheme 6 is rewritten as following:

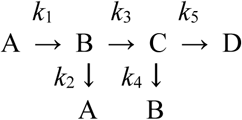

We define the partitions of B that proceeds to C and C that proceeds to D as *p*_1_ and *p*_2_, respectively:

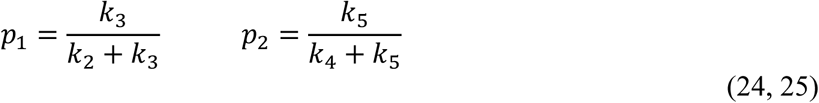

The complete construction of a probability tree diagram for this case is complicated due to the presence of two branching points and not so practical. The application of the recursive use of τ_rxn_ is still feasible. The mean reaction time is the sum of the mean reaction time for three fractions: the fraction of *p*_1_*p*_2_ that proceeds from A to D without any returning, the fraction of *p*_1_(1–*p*_2_) that proceeds from A to C, but returns to B, and the fraction of 1–*p*_1_ that proceeds from A to B, but returns to A (Figure 2C):

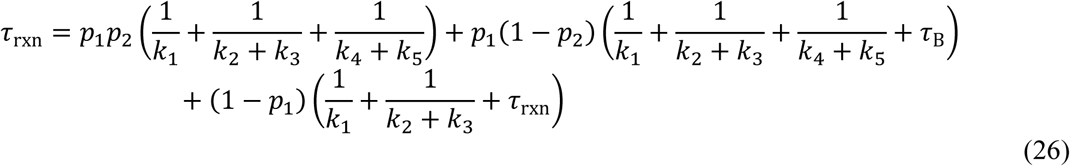

where τ_B_ is the mean reaction time required for B to finish the reaction. Because the conversion of A to B requires the mean reaction time of 1/*k*_1_, τ_rxn_ has the following relationship with τ_B_.

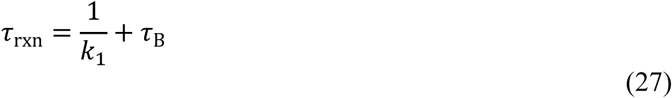

Using this relationship, Eq. 26 is rewritten as:

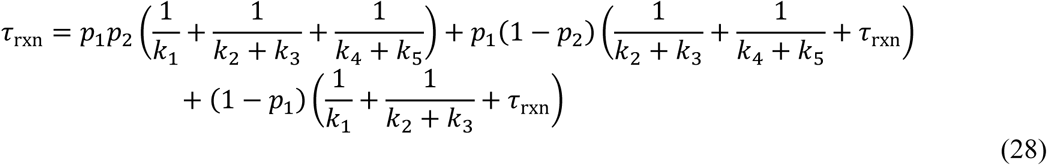

By solving Eq. 28 for τ_rxn_, we obtain the expression of τ_rxn_:

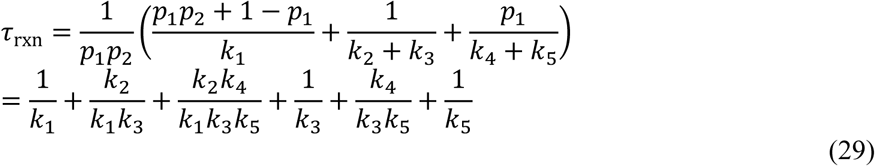

Compared with a reaction with three irreversible steps, the mean reaction time in Eq. 29 has three extra terms: *k*_2_/*k*_1_*k*_3_, *k*_2_*k*_4_/*k*_1_*k*_3_*k*_5_, and *k*_4_/*k*_3_*k*_5_, which are the kinetic costs of the two reversible steps. Apparently, *k*_2_/*k*_1_*k*_3_ and *k*_2_*k*_4_/*k*_1_*k*_3_*k*_5_ are the increase in the total residence time of A due to the first and second reversible steps, respectively, and *k*_4_/*k*_3_*k*_5_ are the increase in the total residence time of B due to the second reversible step. One can verify this easily by setting *k*_2_ or *k*_4_ to 0 and check which extra terms disappear. Again, the mean reaction time is composed of the total residence time of A, B, and C:

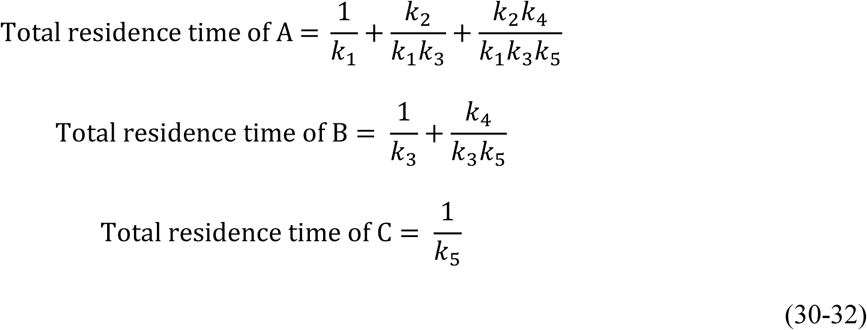

We set the last steps of Schemes 5 and 6 to be irreversible. If all steps are reversible as shown in Scheme 7, it is not possible to define the mean reaction time because the reaction would never reach the completion.

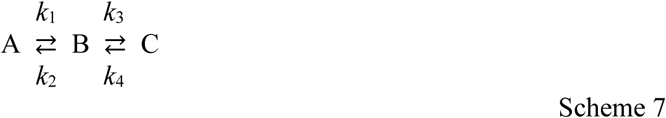

Instead, we determine the mean reaction time for the first arrival to the product by making the last step irreversible, i.e. setting *k*_4_ to 0; therefore, the mean reaction time for the forward reaction (τ_rxn_^f^) of Scheme 7 is same as the mean reaction time for Scheme 5 (Eq. 19).

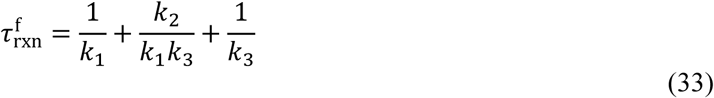

By the same way, we determine the mean reaction time for the reverse reaction (τ_rxn_^r^) of Scheme 7 by setting *k*_1_ to 0.

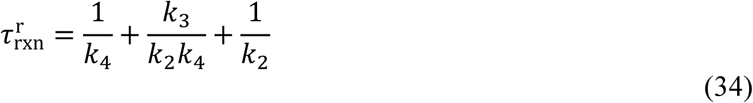

The ratio of τ_rxn_^r^ to τ_rxn_^f^ is the ratio of the time the chemical entity spends in the reverse direction and in the forward direction and identical to the equilibrium constant of the reaction.

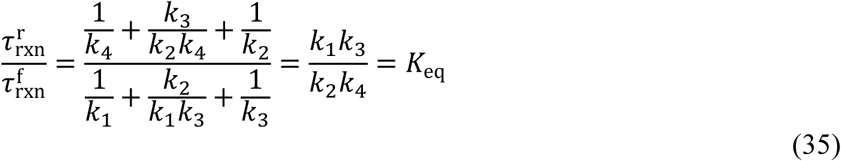

Note that we do not use any approximation, such as steady-state approximation or rapid equilibrium approximation, in deriving this relationship. Whether τ_rxn_^r^ and τ_rxn_^f^ are explicit or implicit, Eq. 35 is valid

## Application of net rate constants

The net rate constant was originally devised to simplify the derivation of kinetic equations for steady-state enzyme-catalyzed reactions.^4^ However, the definition of the net rate constant does not require he steady-state approximation. By designating a net rate constant to a reversible step, we can convert reversible steps to kinetically equivalent irreversible steps, which offers a shortcut to the determination of the mean reaction time. The reaction in Scheme 6 is converted to a reaction with only irreversible steps using net rate constants:

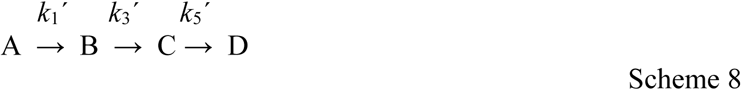

The net rate constant for an irreversible step is identical to the true rate constant of the step; therefore *k*_5_’= *k*_5_. The net rate constant for a reversible step is determined as the forward rate constant multiplied by the fraction of the conversion that eventually finishes the reaction. The fraction is calculated as the ratio of the net rate constant of the subsequent step to the sum of the net rate constant for the subsequent step and the reverse rate constant. The net rate constants are determined by going backward from the last irreversible step as shown below.

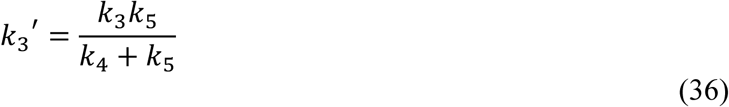

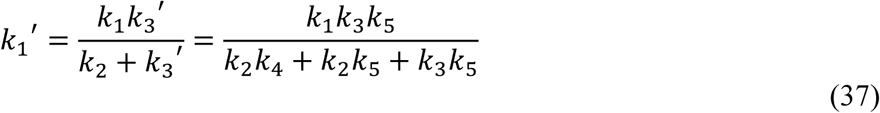

Using Rule 1, we calculate the mean reaction time as:

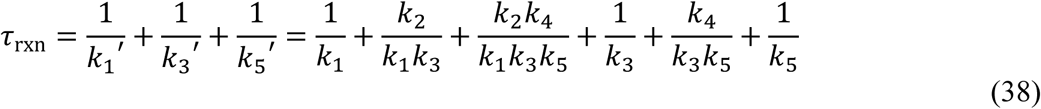

which is identical to Eq. 29. Apparently, the reciprocal of *k*_1_’ is the total residence time of A (Eq. 30), and the reciprocal of *k*_3_’ is the total residence time of B (Eq. 31), and so forth. The reciprocal of the net rate constant for the enzyme-catalyzed reaction was previously called the transit time of the enzyme in the corresponding enzyme form,^6^ which is same as the total residence time we define here for chemical reactions in general. Note that, though Schemes 6 and 8 are kinetically equivalent, and their mean reaction times (and the total residence time at each step) are identical, the reaction trajectories each chemical entity experiences in the two Schemes are distinct.

## Energetic consideration

The reaction energy diagram visually shows the relative magnitude of the energy barrier corresponding to each elementary rate constant of a kinetic model (Figure 3). Using the reaction energy diagram, we can examine how the energy barriers determine each term of the mean reaction time. Figure 3 shows a normative energy diagram for the kinetic model shown in Scheme 6. The height of the barrier for each rate constant corresponds to Δ*G*^‡^, which is linearly proportional to the natural logarithm of the mean lifetime (1/*k*) for each step. The higher the energy barrier is, the longer each step takes to proceed.

**Figure 3.**
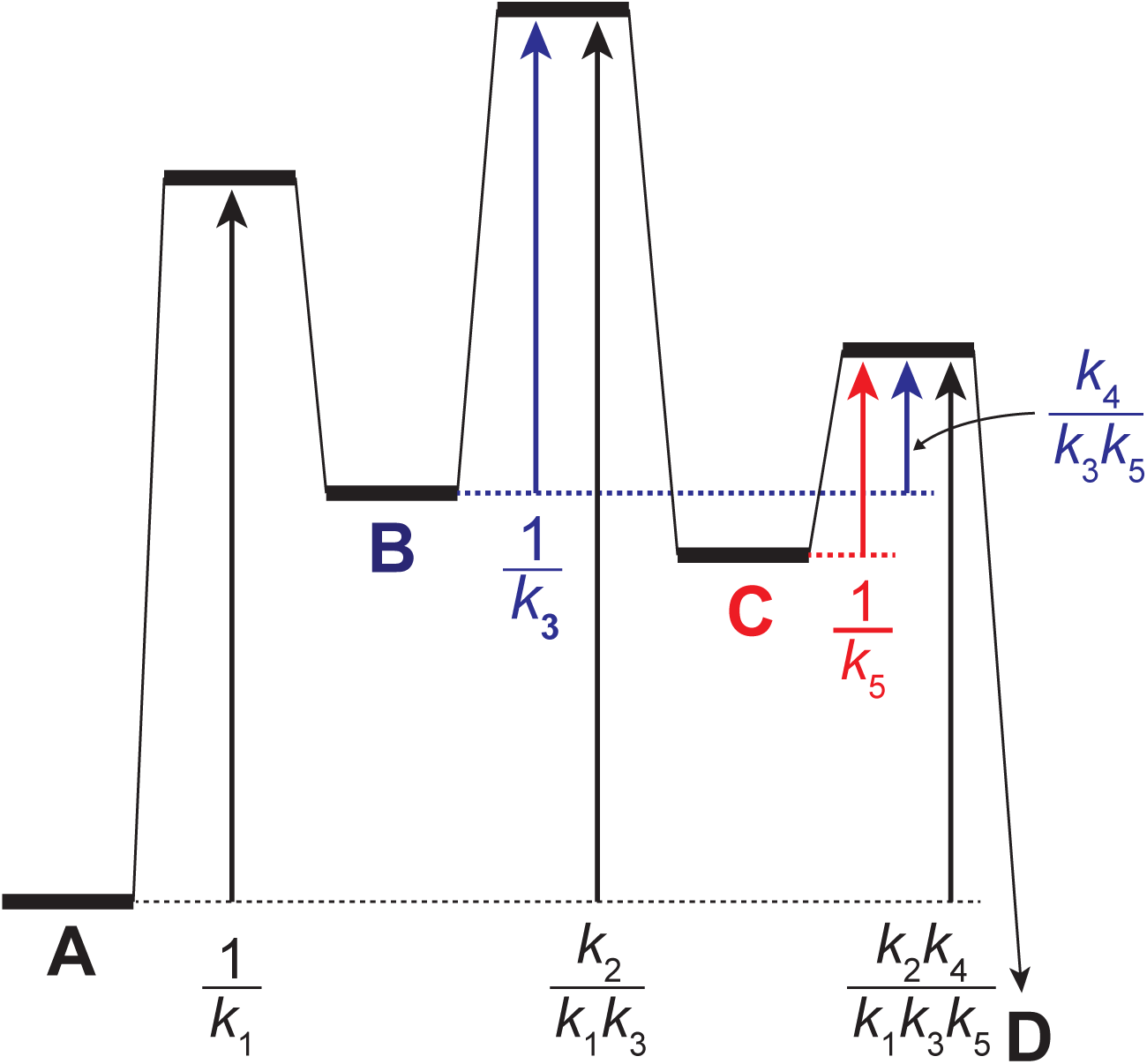
Normative reaction energy diagram for the kinetic model in Scheme 6. Each term of the mean reaction time for the reaction in Scheme 6 is shown as an arrow between the corresponding ground state and transition state. The terms of the total residence times of A, B, and C are indicated by black, blue, and red arrows, respectively.

The expression for the mean reaction time in Eq. 29 has total six terms. Three of them, 1/*k*_1_, 1/*k*_3_, and 1/*k*_5_, corresponds to the energy barriers for A to B, B to C, and C to D, respectively. If the reaction does not have any reversible step, the reaction time will be determined by only these three energy barriers. The rest three terms, *k*_2_/*k*_1_*k*_3_, *k*_2_*k*_4_/*k*_1_*k*_3_*k*_5_, and *k*_4_/*k*_3_*k*_5_, are the kinetic cost of the reversibility of the reaction. The magnitudes of the three terms can be visualized easily on the reaction energy diagram. *k*_2_/*k*_1_*k*_3_ corresponds to the height of the second transition state relative to the energy of A, *k*_2_*k*_4_/*k*_1_*k*_3_*k*_5_ corresponds to the height of the third transition state relative to the energy of A, and *k*_4_/*k*_3_*k*_5_ corresponds to the height of the third transition state relative to the energy of B (Figure 3). Therefore, the total residence time of A (Eq. 30) is determined by the heights of the three subsequent barriers relative to the energy of A. Likewise, the total residence time of B (Eq. 31) is determined by the heights of the two subsequent barriers relative to the energy of B. If the conversion of A to B is irreversible, the energy barrier after B would not affect the total residence time of A. If the conversion of A to B is reversible, however, all the energy barriers till the first irreversible step increase the total residence time of A, and the contribution of each subsequent barrier to the total residence time of A is determined by the height of each barrier relative to the energy of A.

The six terms in Eq. 29 are arranged in Table 1 according to their corresponding ground state (A, B, or C) and transition states (A→B, B→C, and C→D). Each term in the table is the contribution of the corresponding barrier to the total residence time of the corresponding ground state. The pattern of the terms is quite obvious; each term is the ratio of the product of all the reverse rate constants to the product of all the forward rate constants between the ground state and the transition state. Note that the reverse rate constant corresponding to the transition state is not included. The sum of the terms in each row corresponds to the total residence time of each ground state. The sum of the terms in each column is the contribution of each transition state to the mean reaction time. The diagonal of the table contains the mean lifetime of each ground state, which is also the residence time of each ground state in case all the steps are irreversible. Above the diagonal are the kinetic costs of the reversibility, i.e. the increase in the residence time of each ground state due to the barriers connected through reversible steps.

**Table 1.** Terms of the mean reaction time. The terms of the mean reaction time for the kinetic model in Scheme 6 (Eq. 29) are arranged according to the corresponding ground state and the transition state. The total residence time is the sum of terms in each row (Eqs. 30-32).

| Ground states | Transition states |  |  | Total residence times |
| --- | --- | --- | --- | --- |
|  | A→B | B→C | C→D |  |
| A | $\frac{1}{k_1}$ | $\frac{k_2}{k_1 k_3}$ | $\frac{k_2 k_4}{k_1 k_3 k_5}$ | $\frac{1}{k_1} + \frac{k_2}{k_1 k_3} + \frac{k_2 k_4}{k_1 k_3 k_5}$ |
| B | 0 | $\frac{1}{k_3}$ | $\frac{k_4}{k_3 k_5}$ | $\frac{1}{k_3} + \frac{k_4}{k_3 k_5}$ |
| C | 0 | 0 | $\frac{1}{k_5}$ | $\frac{1}{k_5}$ |

## Rate-limiting step versus most time-consuming process

The analysis of the terms comprising the mean reaction time provides us with a new perspective on the issue of the rate-limiting step. Identifying the rate-limiting step of a complex reaction is a common practice in chemistry. The rate-limiting step is frequently compared to a bottle neck, implying that the step with the smallest rate constant is rate-limiting. This definition of the rate-limiting step works fine with the complex reaction only with *irreversible* steps, where the contribution of each step to the mean reaction time is independent and separable (the diagonal terms in Table 1). For the complex reaction with *reversible* steps, however, this definition of the rate-limiting step does not work. For example, in case of the reaction depicted in Figure 3, *k*_1_ is the smallest rate constant out of the three forward rate constants (*k*_1_, *k*_3_, and *k*_5_). However, the greatest contributor to the mean reaction time is not 1/*k*_1_ but *k*_2_/*k*_1_*k*_3_. Moreover, *k*_2_/*k*_1_*k*_3_ corresponds to not a barrier of a single step but the height of the second transition state relative to the energy level of A. This difficulty in defining the rate-limiting step for the complex reaction with reversible steps (especially enzyme-catalyzed reactions) has been addressed before.^7, 8^ The step with the smallest rate constant (the most difficult step) or the step that accumulates the most intermediate (the least conducive step) is not necessarily rate-limiting. The step with the highest transition state is not always rate-limiting, either. In conclusion, for the complex reaction with *reversible* steps, identifying a *single* step that limits the overall reaction rate is improper in many cases. The off-diagonal terms in Table 1 clearly show the reason for this conclusion. The contribution of the steps to the mean reaction time are complex. The transition state of a single step affects the residence time of all preceding ground states connected by reversible steps. Likewise, the residence time of a ground state is affected by all subsequent transition states connected by reversible steps and the transition state of the first irreversible step.

The easily identifiable and well-defined terms in Table 1 allows us to identify the *most time-consuming* process (not a step) instead of the rate-limiting step improper for the complex reactions with reversible steps. The most time-consuming process is the largest term of the polynomial expression of the mean reaction time (Table 1). The most time-consuming process corresponds to the pair of a ground state and one of the subsequent transition states that has the largest energy gap. In Figure 3, *k*_2_/*k*_1_*k*_3_ has the largest energy gap (longest arrow), which is between A and the second transition state, and the most time-consuming process of the reaction is that A crosses the second transition state. In enzyme-catalyzed reactions, it is quite valuable to identify the step most sensitive to the isotope substitution of the substrate. Assuming that the isotope substitution only affects the levels of the transition state (no equilibrium isotope effect), the *most sensitive step*,^7^ which was also called the *kinetically significant barrier*,^9^ is the transition state that contributes most to the mean reaction time. The sum of each column in Table 1 corresponds to the contribution of each transition state to the mean reaction time. Therefore, the transition state that has the largest sum corresponds to the most sensitive step. The *least conducive step*^7^ that accumulates the most intermediate is also sometimes informative. The sum of each raw in Table 1 corresponds to the total residence time of each ground state. The subsequent step of the ground state with the greatest total residence time (also the smallest net rate constant), which was also called the *kinetically significant intermediate*,^9^ therefore, is the least conducive step. Note that the *most-time consuming process* is likely to occur between the *kinetically significant intermediate* and the *kinetically significant barrier*, but this is not always the case.

## Generalization of the formula for the mean reaction time

Now we derive the expression for the mean reaction time for a generalized kinetic model with *n* intermediates (I_1_, I_2_, …, and I_n_) shown in Scheme 9.

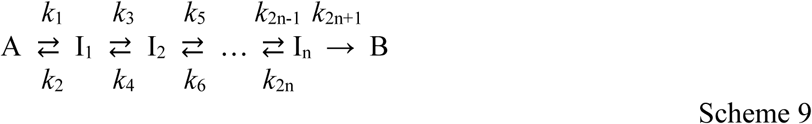

As shown in Table 1, the total residence time of a ground state is the sum of the residence time imposed by all the subsequent transition states. Therefore, we write the formula for the mean reaction time as the sum of the total residence time of all ground states:

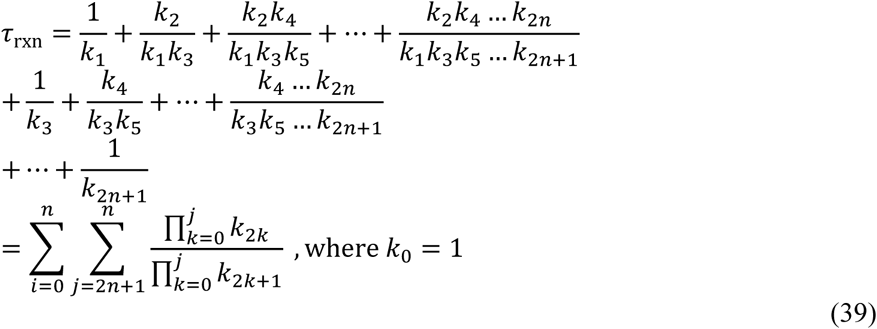

We can arrange the terms in Eq. 39 according to the corresponding ground state and transition state as in Table 1. The diagonal terms are the mean lifetime of the ground state (1/*k*_1_, 1/*k*_3_, 1/*k*_5,_ and so forth), and the off-diagonal terms are the kinetic cost of the reversibility. As we discussed above, the sum of each row is the total residence time of each ground state, and the sum of each column is the contribution of each transition state to the mean reaction time.

The generalized formula (Eq. 39) is also applicable to any model that contains one or more irreversible steps in the middle of the kinetic model. The mean reaction time for the model with irreversible steps can be obtained by setting the rate constants for the reverse reaction at the irreversible steps to 0. For example, the conversion of I_k-1_ and I_k_ is irreversible, we set *k*_2k_ to 0, which removes any term that contains *k*_2k_ in Eq. 39. Note that the rate constants for the reverse reactions have indexes of even numbers and always appear in the numerators in Eq. 39. Also, an irreversible step in the middle of the reaction breaks the reaction into kinetic segments, and the mean reaction time is the sum of the mean reaction time of each segment. The reaction in Scheme 10 has an irreversible step between B and C:

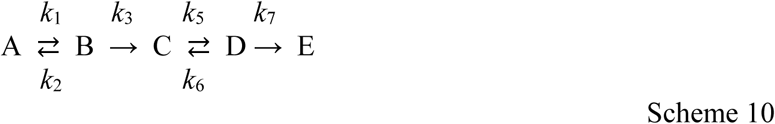

The irreversible step breaks the reaction to two kinetic segments: A ⇄B → C and C ⇄D → E. Because each segment is same as the reaction in Scheme 5, the mean reaction time for the first segment is 1/*k*_1_ + *k*_2_/*k*_1_*k*_3_ + 1/*k*_3_, and that for the second segment is 1/*k*_5_ + *k*_6_/*k*_5_*k*_7_ + 1/*k*_7_. Therefore, the mean reaction time for the whole reaction is:

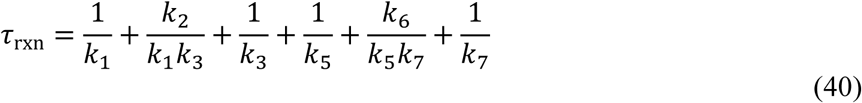

One can also derive Eq. 40 by setting *k*_4_ to 0 in Eq. 39.

## Application to enzyme kinetics

Now we apply the mean reaction time to enzyme kinetics. Enzyme kinetics have been described with the reaction time instead of the rate.^10, 11^ The reciprocal form of the rate equation of the enzyme-catalyzed reaction shows that the reaction time is expressed as the sum of terms dependent on and independent of the concentrations of the substrates. Also, it has been shown that the reciprocal of the net rate constant corresponds to the transit time of each enzyme species (e.g. E, ES, EP).^6^ However, the expression of the reaction time was obtained only after the rate equation was derived. Here, we derive the expression of the mean reaction time directly from the kinetic model without deriving the complex rate equation based on the steady-state approximation.

To calculate the mean reaction time for the enzyme-catalyzed reaction such as Scheme 1, we need to address two issues. First, the enzyme-catalyzed reaction starts with a bimolecular reaction, the association of E and S, and *k*_1_ is a second-order rate constant. The total residence time of the free enzyme (E) cannot be calculated only with *k*_1_ because the residence time is dependent on the concentration of the substrate. Instead of *k*_1_, we need to use a pseudo-first-order rate constant, *k*_1_[S], which is a common practice in enzyme kinetics. Second, the enzyme-catalyzed reaction is a cyclic reaction, which starts with E and ends with E. As E is regenerated after each cycle (i.e. E is catalytic), we cannot define the mean reaction time to be the time required for the completion of the reaction. We address this issue by setting up a theoretical kinetic model for a single turnover, where E is converted to E^*^ after finishing one catalytic turnover, and the mean reaction time is the time required for the single turnover. With these modifications, the reaction in Scheme 1 is rewritten as:

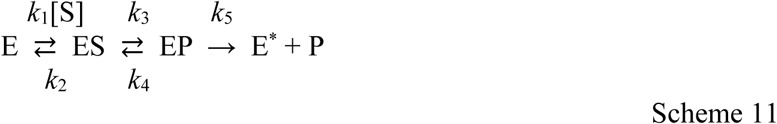

Now the reaction is kinetically equivalent to the reaction in Scheme 6, and the mean reaction time is:

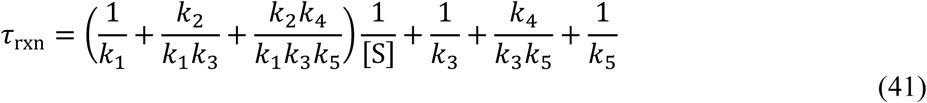

Eq. 41 shows that the coefficient of 1/[S] and the remaining term are actually the reciprocal forms of *k*_cat_/*K*_M_ and *k*_cat_ in Eqs. 6 and 7.

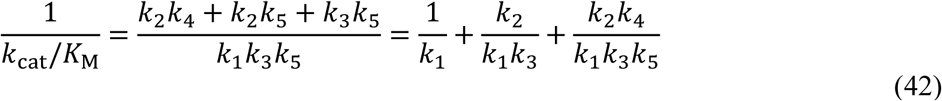

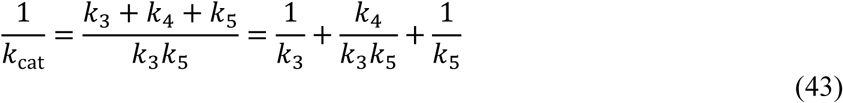

Therefore, Eq. 41 can be expressed using *k*_cat_/*K*_M_ and *k*_cat_ as:

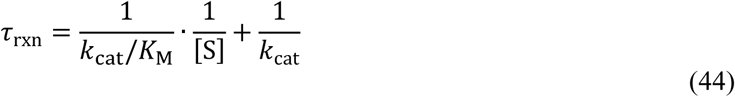

Eqs. 41 and 44 demonstrate that we can derive the expressions for *k*_cat_ and *k*_cat_/*K*_M_ as functions of elementary rate constants simply by calculating the mean reaction time of an enzyme-catalyzed reaction even without using the steady-state approximation. Note that the only approximation we employ here is the use of the pseudo-first-order rate constant. Also, Eq. 41 gives us a valuable insight on the meanings of *k*_cat_ and *k*_cat_/*K*_M_. 1/*k*_cat_ is the sum of the total residence time of all bound forms of E (ES and EP in Scheme 11) for the single turnover, 1/(*k*_cat_/*K*_M_[S]) is the total residence time of the free form of E for the single turnover.

When the enzyme-catalyzed reaction reaches the steady state, we can assume that the concentrations of all the enzyme forms (free or bound) do not change, and the reaction rate is constant. Under this condition, the mean reaction time is explicit so that 1/τ_rxn_ is the rate constant of the reaction. One enzyme converts one substrate molecule to a product during the single turnover, and one enzyme can generate 1/τ_rxn_ product molecules per unit time. Therefore, the total concentration of the product produced per unit time, which is *v* by definition, is equal to [E_t_]/τ_rxn_. Using this relationship, we can rewrite Eq. 44 as:

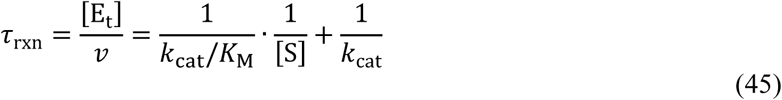

which is indeed the reciprocal form of the Michaelis–Menten equation. The reciprocal form of the Michaelis–Menten equation has been used not only to fit the experimental data (e.g. Lineweaver–Burk plot) but also for analyzing the enzyme kinetics in the time domain.^10, 11^ Here, we derive the same formula only by calculating the mean reaction time. The generalized approach to determine the mean reaction time, therefore, is surprisingly simple shortcut to derive the expressions for *k*_cat_/*K*_M_ and *k*_cat_. which is indeed the reciprocal form of the Michaelis–Menten equation. The reciprocal form of the Michaelis–Menten equation has been used not only to fit the experimental data (e.g. Lineweaver–Burk plot) but also for analyzing the enzyme kinetics in the time domain.^10, 11^ Here, we derive the same formula only by calculating the mean reaction time. The generalized approach to determine the mean reaction time, therefore, is surprisingly simple shortcut to derive the expressions for *k*_cat_/*K*_M_ and *k*_cat_.

Here we demonstrate the use of the mean reaction time using a simple kinetic model with only one substrate and one product (Scheme 1). The method can be applied easily to more complex kinetic models with multiple substrates and products. Deriving the formula of the mean reaction time for the kinetic model with branched pathways is possible, but somewhat complex. However, the true value of the mean reaction time is not as a shortcut to derive a rate equation but as a mathematical tool to demonstrate how the individual steps and their elementary rate constants contribute to the reaction rate. The arrangement of the terms of the mean reaction time as shown in Table 1 offers a surprisingly simple way to visualize and dissect the contribution of each ground state and transition state to the kinetics of the reaction.

